# A structural variant in the 5’-flanking region of the TWIST2 gene affects melanocyte development in belted cattle

**DOI:** 10.1101/077065

**Authors:** Nivedita Awasthi Mishra, Cord Drögemüller, Vidhya Jagannathan, Rémy Bruggmann, Julia Beck, Ekkehard Schütz, Bertram Brenig, Steffi Demmel, Simon Moser, Heidi Signer-Hasler, Aldona Pieńkowska-Schelling, Claude Schelling, Ronald Rongen, Stefan Rieder, Robert N. Kelsh, Nadia Mercader, Tosso Leeb

**Affiliations:** Institute of Genetics, Vetsuisse Faculty, University of Bern, 3001 Bern, Switzerland; DermFocus, Vetsuisse Faculty, University of Bern, 3001 Bern, Switzerland.; Swiss Competence Center of Animal Breeding and Genetics, University of Bern, Bern University of Applied Sciences HAFL & Agroscope, 3001 Bern, Switzerland; Interfaculty Bioinformatics Unit, University of Bern, 3012 Bern, Switzerland; Chronix Biomedical, 37073 Göttingen, Germany; Department of Molecular Biology of Livestock, Georg August University, 37077 Göttingen, Germany; Bern University of Applied Sciences, School of Agricultural, Forest and Food Sciences, 3052 Zollikofen, Switzerland; Clinic for Reproductive Medicine, University of Zurich, 8057 Zurich, Switzerland; Dutch Belted Cattle Association, 7211 EM Eefde, The Netherlands; Agroscope, Swiss National Stud Farm, 1580 Avenches, Switzerland; Department of Biology and Biochemistry, University of Bath, Claverton Down, Bath BA2 7AY, United Kingdom; Institute of Anatomy, University of Bern, 3012 Bern, Switzerland

## Abstract

Belted cattle have a circular belt of unpigmented hair and skin around their midsection. The belt is inherited as a monogenic autosomal dominant trait. We mapped the causative variant to a 54 kb segment on bovine chromosome 3. Whole genome sequence data of 2 belted and 130 control cattle yielded only one private genetic variant in the critical interval in the two belted animals. The belt-associated variant was a copy number variant (CNV) involving the quadruplication of a 6 kb non-coding sequence located approximately 16 kb upstream of the *TWIST2* gene. Increased copy numbers at this CNV were strongly associated with the belt phenotype in a cohort of 239 cases and 1303 controls (p = 1.3 x 10^-278^). We hypothesized that the CNV causes aberrant expression of *TWIST2* during neural crest development, which might negatively affect melanoblasts. Functional studies showed that ectopic expression of bovine *TWIST2* in neural crest in transgenic zebrafish led to a decrease in melanocyte numbers. Our results thus implicate an unsuspected involvement of TWIST2 in regulating pigmentation and reveal a non-coding CNV underlying a captivating Mendelian character.

**Author Summary:** Belted cattle, a spontaneous coat color mutant, have been recognized at least 600 years ago. The striking pigmentation pattern probably has arisen in medieval cattle of the Alpine region. The belt still segregates in Brown Swiss cattle and it has become a breed-defining character in the Lakenvelder or Dutch Belted cattle. The belted allele has also been introgressed into Galloways to form the Belted Galloways. We report here the causative genetic variant, a non-coding copy number variant (CNV) upstream of the *TWIST2* gene. We hypothesize that the CNV leads to ectopic expression of TWIST2 in the neural crest, which negatively affects melanocyte development. Overexpression of bovine TWIST2 in transgenic zebrafish embryos led to a decrease in melanocyte numbers, which provides functional support for our hypothesis.

## Introduction

Coat color has been a long-standing model trait for geneticists due to the ease of phenotype recording. Coat color depends on the presence of pigment cells. In mammals, melanin-synthesizing melanocytes, which occupy the skin and hair follicles throughout the entire body surface, are responsible for pigmentation [1]. During embryonic development melanocytes are formed from melanoblasts, which originate in the neural crest and migrate through the developing embryo in order to reach their final position on the body [2]. This developmental program requires a delicate level of regulation to ensure that the correct number of cells reaches their final destination [3–5]. An over-proliferation of cells that have left their surrounding tissue leads to naevi, and in rare cases to congenital melanomas which lead to a poor prognosis [6–8]. In contrast, if too few melanoblasts are produced or survive, this will lead to partially or completely unpigmented phenotypes, including so-called “white-spotting” phenotypes characterized by patches of unpigmented skin and/or hair [9]. Domestic animals with such hypopigmented phenotypes have been highly valued due to their striking appearance and have often been actively selected in animal breeding. Modern domestic animals thus constitute a unique reservoir of spontaneous pigmentation mutants. Their favorable population structure facilitates the discovery of functional genetic variation including some interesting non-coding structural variants with regulatory effects [10–15].

A symmetrical belt of unpigmented skin circling the midsection has been observed in several mammalian species. It is thought that these belted phenotypes are due to downregulated melanoblast formation or early melanoblast losses in neural crest development. Belted mice have sequence variants in the *Adamts20* gene encoding a secreted metalloprotease [16], which was shown to be required for melanoblast survival [17]. A complex structural rearrangement involving duplication of the *KIT* gene was identified in belted pigs, whose belt includes the forelimbs and is localized more cranially than the one in *Adamts20* mutant mice [18,19].

Belted cattle exist in at least three different breeds: Brown Swiss, (Belted) Galloway, and Lakenvelder, which are also known as Dutch Belted. The belt in these cattle breeds has an intermediate position; it is more caudal than the belt in pigs and more cranial than the belt in *Adamts20* mutant mice (Figure 1). The belt in cattle is inherited as a monogenic autosomal dominant trait and linked to the telomeric end of chromosome 3, which does not contain any known coat color gene [20,21]. The objective of this study was to identify the causative genetic variant and to provide a hypothesis on the functional mechanism leading to the partial lack of melanocytes in belted cattle.

**Figure 1.**
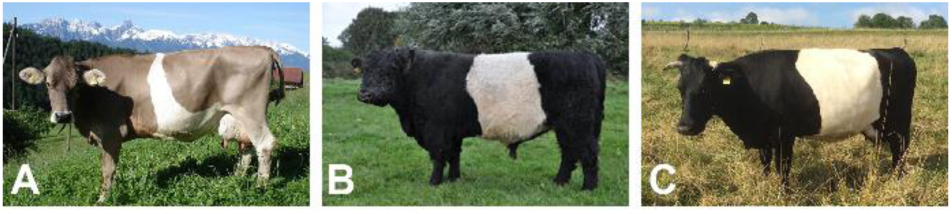
Belted phenotype in different cattle breeds. (A) Brown Swiss. (B) Belted Galloway. (C) Lakenvelder or Dutch Belted. The width and shape of the belt is variable and in some belted animals the unpigmented area does not fully circle the animal.

## Results

### Positional cloning of the belt sequence variant reveals a CNV upstream of *TWIST2*

We had previously shown that the belt locus maps to a 336 kb interval at the telomeric end of chromosome 3 and that belted animals from the Brown Swiss, Belted Galloway and Lakenvelder breeds share the same haplotype in the critical interval [21]. In order to further refine this interval we now genotyped 116 belted cattle on the Illumina bovine high density beadchip, phased the genotype data and searched for a shared haplotype. All belted animals had at least one copy of a shared haplotype spanning 22 consecutive markers (Table S1). Thus, the refined boundaries of the new critical interval are defined by the flanking markers on either side of this shared haplotype block. The refined critical interval consisted of only 54 kb and spanned from position 118,578,893 to 118,633,153 on chromosome 3 (Figure 2A).

**Figure 2.**
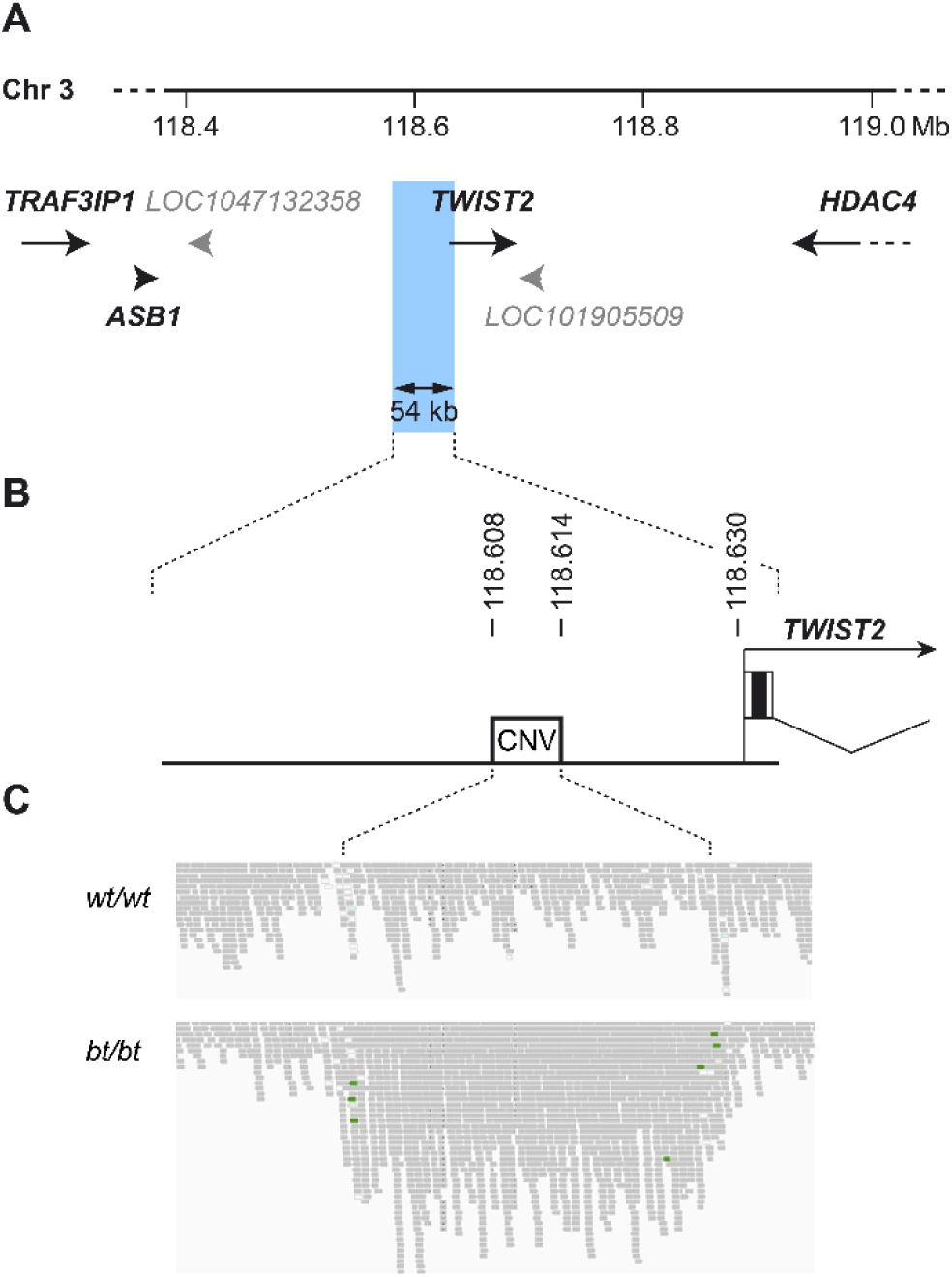
Genomic context of the belt locus. (A) Haplotype analysis defined a critical interval of 54 kb indicated by blue shading for the belt causative variant (Chr3:118,578,893-118,633,153, UMD3.1 assembly). The belt locus mapped to a gene poor region containing *TWIST2* as the only known gene (NCBI annotation release 105; protein coding genes are shown in black, predicted non-coding RNA genes in grey). (B) A 6 kb CNV is located within the critical interval and approximately 16 kb upstream of the transcription start site of *TWIST2*. (C) Experimental identification of the CNV. IGV screenshots of the illumina short read sequences illustrate a ~4-fold increased coverage in a homozygous belted (*bt/bt*) cattle with respect to a control animal (*wt/wt*) and several read-pairs with incorrect read-pair orientation at the boundaries of the CNV (indicated in green).

For the identification of potentially causative genetic variants, we sequenced the genomes of two homozygous belted cattle and compared the data to 130 genomes from control cattle. A standard bioinformatic pipeline for the detection of single nucleotide variants or small indels did not reveal any private homozygous variants in the two cases in the critical interval. We therefore visually inspected the short-read sequencing data in the critical interval and identified an amplification of a 6 kb sequence region (Figure 2B,C). Both belted animals had a ~4-fold increased read coverage with respect to control animals. The 6 kb CNV was located approximately 16 kb upstream of the *TWIST2* gene. We did not detect any other structural variant in the critical interval.

We designed a digital droplet PCR (ddPCR) assay to determine the copy number of the CNV in a large cohort of 239 belted cattle and 1303 non-belted cattle. The observed copy numbers varied between 2 and 12 with the most frequently observed values being 2, 5 and 8, which most likely corresponded to genotypes *1/1*, *1/4* and *4/4*. The belt phenotype was strongly associated with an increased copy number at the CNV (p = 1.3 x 10^-278^, Fisher’s exact test; Table 1).

**Table 1.** Association of the belt phenotype with the CNV.

| Copy number | Belted cattle<br>(N = 239) | Non-belted cattle<br>(N = 1303) |
| --- | --- | --- |
| 2 | 2<sup>a</sup> | 1301 |
| 3 | 2 | - |
| 4 | 12 | - |
| 5 | 62 | 2<sup>a</sup> |
| 6 | 13 | - |
| 7 | 42 | - |
| 8 | 96 | - |
| 9 | 9 | - |
| 12 | 1 | - |

<sup>a</sup>We observed a nearly perfect association between increased copy number and the belted phenotype. The four discordant animals in this analysis may in fact have been incorrectly phenotyped (see Materials and Methods).

### Does the CNV affect TWIST2 expression?

TWIST2 is normally expressed in the skin and craniofacial cartilages, but not in the neural crest [22,23]. We performed an RNA-seq experiment to quantify *TWIST2* RNA expression in adult skin from belted and non-belted cattle, but found no significant difference (Table S2). The strictly dominant mode of inheritance of the belt suggests a gain of function of the *bt* allele. Given that the boundaries of the unpigmented area in belted animals are very even and mostly bilaterally symmetrical, this gain of function might supposedly take place during early melanoblast development. We hypothesized that increased copy numbers at the CNV might lead to an ectopic expression of TWIST2 during neural crest development. Accordingly, ectopically expressed TWIST2 might decrease the number of melanoblasts and thus lead to the unpigmented skin area of the belt. Unfortunately, it is currently not possible to test this hypothesis by directly quantifying TWIST2 expression in the developing neural crest of bovine embryos.

### Functional analysis of TWIST2 overexpression

As bovine embryos are not readily accessible for experimental research, we investigated the effect of TWIST2 overexpression in transgenic zebrafish embryos. We prepared expression constructs with a zebrafish *mitfa* promoter fragment that has been shown to drive expression in premigratory neural crest cells and differentiating melanocytes [24,25]. The control construct pmitfa_EGFP contained the coding sequence of enhanced green fluorescent protein (EGFP) as a reporter under the control of the *mitfa* promoter. Additionally we prepared a second construct pmitfa_btaTWIST2_EGFP, which additionally contained the coding sequence for bovine TWIST2 separated from the EGFP coding region by an internal ribosome entry site (IRES) so that TWIST2 overexpressing cells were expected to be distinguishable in transient transgenic embryos by their EGFP expression.

We injected both constructs into fertilized zebrafish eggs and analyzed the expression of EGFP at 35 hours post fertilization (hpf) in four replicate experiments. About 80% of more than 100 surviving zebrafish embryos injected with the control construct pmitfa_EGFP showed several EGFP-positive cells in a pattern resembling developing melanoblasts (Figure 3A,C,D). In contrast, we never observed any EGFP-positive cell in the zebrafish embryos injected with pmitfa_btaTWIST2_EGFP (Figure 3B). For maximum sensitivity we confirmed these results by immunostaining the transgenic zebrafish embryos with an anti-GFP antibody. We again detected EGFP expression only in the control embryos, but not in embryos that were injected with pmitfa_btaTWIST2_EGFP (Figure 3E-J). We additionally stained for the expression of sox10 to investigate whether there were any gross changes to neural crest development. The comparison of sox10 expression patterns did not indicate any major differences between the control embryos and the embryos overexpressing TWIST2 (data not shown).

**Figure 3.**
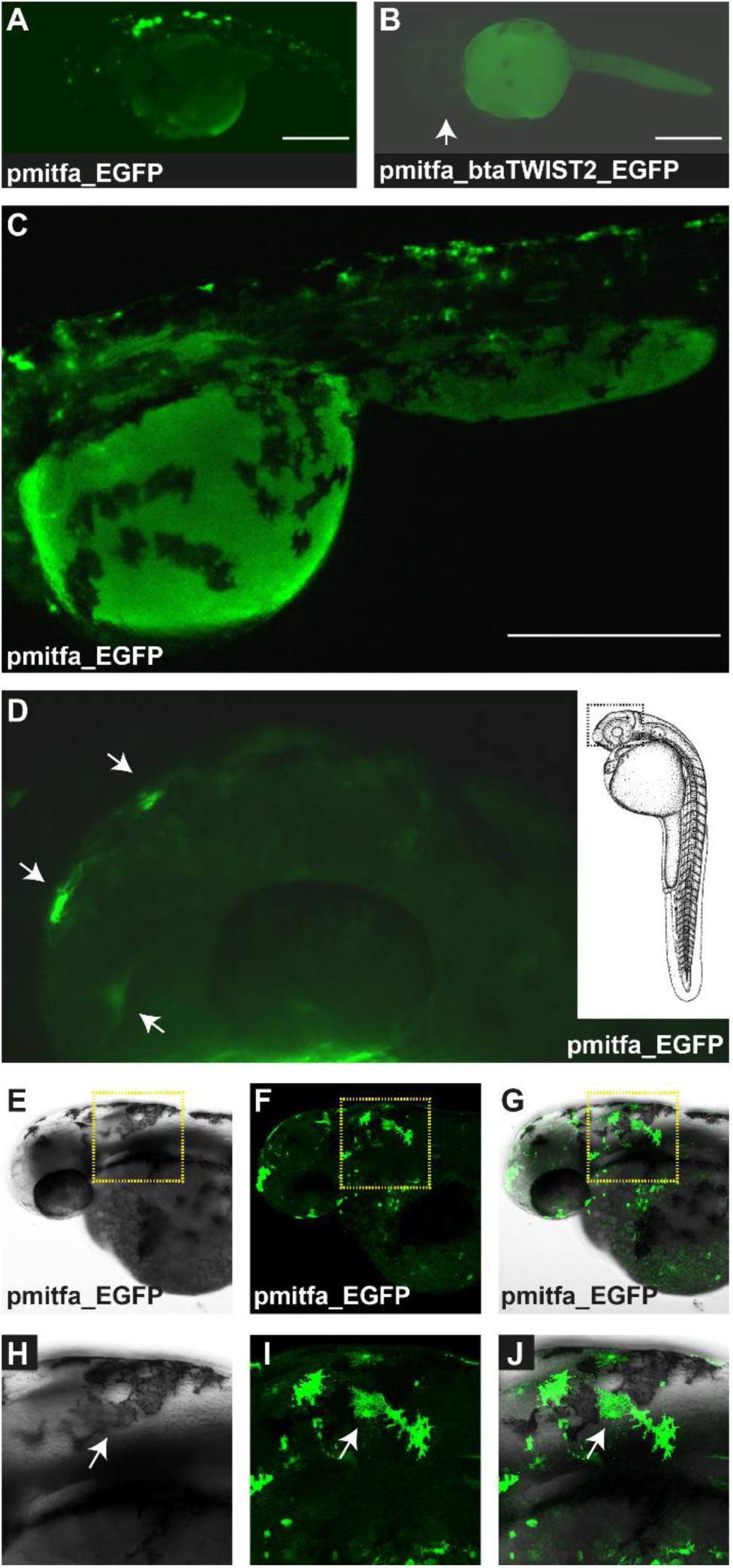
Effect of TWIST2 expression on melanoblasts. (A) A zebrafish 35 hpf embryo injected with the control construct pmitfa_EGFP, which drives expression of green fluorescent protein under the control of the zebrafish mitfa promoter. The head is to the left. Individual EGFP positive cells can be seen. (B) In a zebrafish injected with the construct pmitfa_btaTWIST2_EGFP, driving the bicistronic expression of bovine TWIST2 and EGFP, no EGFP-positive cells can be seen. The head is also pointing to the left. The arrow indicates the position of the eye. The green signal was enhanced for maximum sensitivity. Therefore, the yolk shows some green autofluorescence, which lacks the punctuate pattern as seen in individual cells. (C) The trunk region of another control embryo at higher magnification. The head is to the left. Multiple EGFP-positive cells can be seen. (D) Head of a control embryo. The dashed square in the sketch on the right illustrates the anatomical structures. Three individual EGFP-positive cells can be seen (arrows). (E-H) All pictures show the same embryo injected with the control construct. The area indicated by the yellow dashed square in E-G is magnified in panels H-J. The brightfield images E and H show individual darkly pigmented melanocytes. Images F and I show the green immunofluorescence of an anti-EGFP antibody. G and J show the merged brightfield and green fluorescent images. An individual EGFP-positive melanocyte is indicated by arrows. The scale bars in panels A, B, and C correspond to 1 mm.

We hypothesized that the lack of visible EGFP expression in the zebrafish injected with pmitfa_btaTWIST2_EGFP was due to a negative regulatory effect of bovine TWIST2 on melanoblasts. Our experimental strategy was expected to lead to mosaic animals, where only a fraction of the cells integrated the expression casettes into the genome. In an additional experiment we quantified the number of visible melanocytes in TWIST2 overexpressing versus control embryos. We observed a significant reduction of melanocytes in the head region (p < 0.001). In the trunk we also consistently observed lower mean melanocyte numbers in the TWIST2 transgenic embryos. However, as the variance of melanocyte numbers in the trunk was much bigger than in the head region, this difference was significant only in one out of three replicate epxeriments (Figure 4).

**Figure 4.**
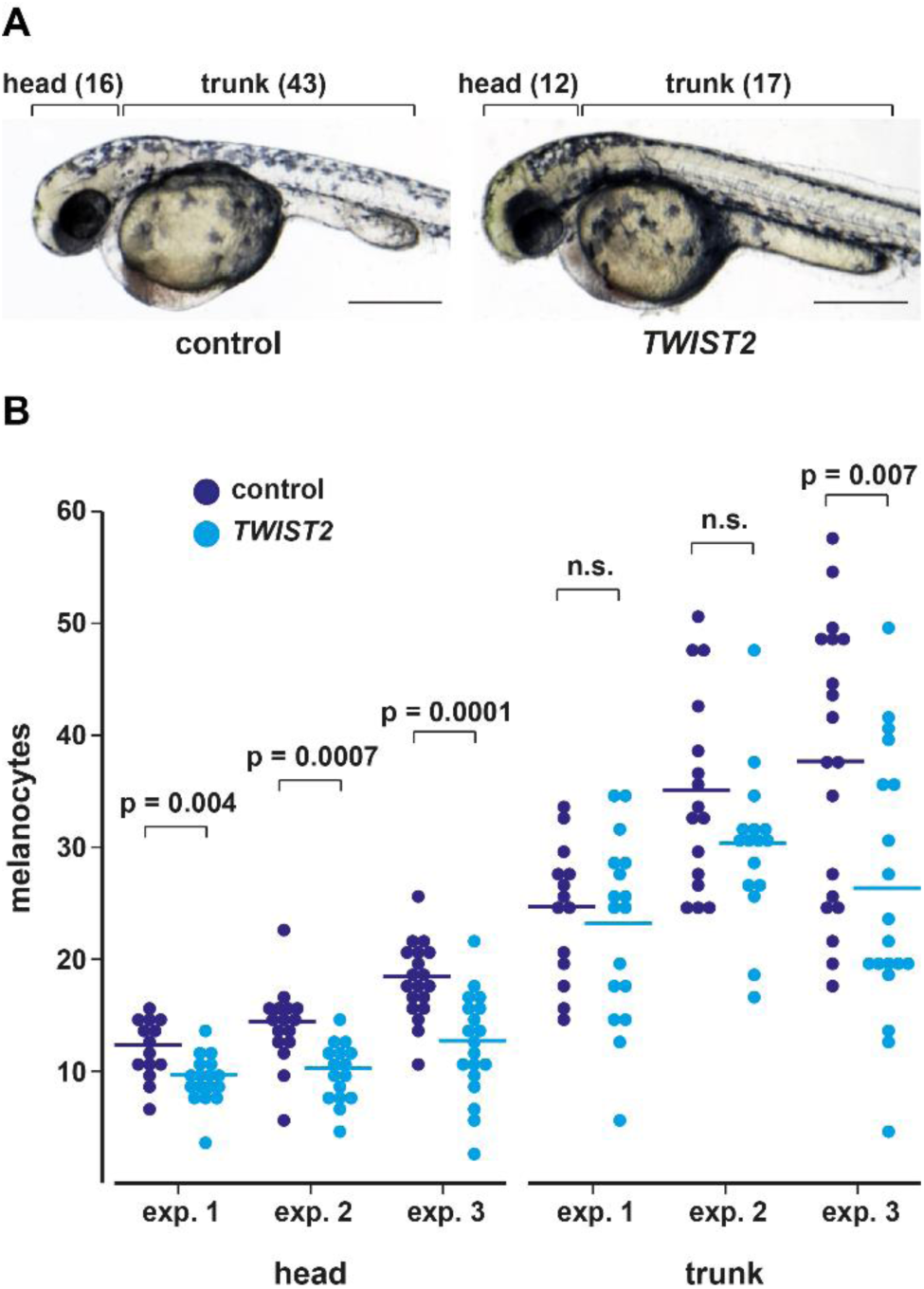
Reduction of melanocytes in zebrafish embryos expressing bovine TWIST2 at 35 hpf. (A) Representative images of zebrafish embryos injected with either a control construct (pmitfa_EGFP) or a construct driving the expression of bovine TWIST2 under control of the zebrafish *mitfa* promoter (pmitfa_btaTWIST2_EGFP). Melanocyte counts for head and trunk are given in brackets. Scale bars correspond to 1 mm. (B) Melanocytes in the head and trunk were counted in three experiments (controls: n = 14, 16, 19; TWIST2: n = 17, 16, 19). The TWIST2 overexpressing embryos had significantly fewer head melanocytes. A similar trend was visible in the trunk.

## Discussion

In this study, we identified the amplification of a 6 kb genomic sequence in the 5’-flanking region of the *TWIST2* gene as most likely causative variant for the belted phenotype in cattle. The haplotype analysis greatly refined the critical interval to only 54 kb. The small size of the shared haplotype is consistent with a relatively old origin of the *bt* allele, having arisen before the separation of modern cattle breeds. A 15^th^ century painting from Austria already depicts a belted ox suggesting that this allele is at least 600 years or roughly 200 generations old (Figure S1). The haplotypes in belted Brown Swiss cattle were more diverse than in Lakenvelder or Belted Galloway. Thus the genetic data point to an origin of the *bt* allele in the alpine region. Historic records are in agreement with an alpine origin of the *bt* allele and indicate that the *bt* allele may subsequently have spread north-west and was introgressed into Dutch cattle to eventually form the Dutch Belted Cattle and into Scottish cattle to eventually form the Belted Galloways [26].

The precise fine-mapping facilitated the search for candidate causative variants. The 54 kb critical interval did not contain any gaps in the reference genome assembly. Thus, our Illumina re-sequencing approach had a good probability to really unravel all sequence variants in this relatively small region. There are only 483 coding nucleotides within the critical interval and these encode the entire 160 amino acid bovine TWIST2 protein. We can reliably exclude the possibility of any coding variants being the cause of the belted phenotype. The CNV was the best associated variant in the critical interval and showed the expected genotypes in 1536 out of 1540 genotyped cattle (99.7%). As the phenotype assignment in the four discrepant animals was solely based on their identity documents, we assume that in these 4 animals the phenotypes were incorrectly assigned (see Materials and Methods). Another potential explanation for the imperfect association could be additional mutation events on the *bt*-associated haplotype that affect the ddPCR genotyping method (e.g. in one of the primer binding sites).

The strictly dominant mode of inheritance of the belt phenotype suggested a gain-of-function allele, such as e.g. an allele with a non-coding regulatory variant leading to ectopic expression of a gene. *TWIST2* represents an obvious candidate gene for such a potentially ectopically expressed gene. It is the only annotated protein-coding gene within 250 kb of the critical interval and the belt-associated CNV is actually located in the 5’-flanking region of *TWIST2*. TWIST2 is a basic helix loop helix (bHLH) transcription factor with a postulated role in early development influencing chondrogenic and dermal tissues [23,27]. *TWIST2* loss-of-function variants in humans cause several complex malformation disorders: ablepharon-macrostomia syndrome (AMS, MIM 200110), Barber-Say syndrome (BSS, MIM 209885) and Setleis syndrome or focal facial dermal dysplasia type III (FFDD3, MIM 227260), all being ectodermal dysplasias with complex facial malformations [28–30]. In one Setleis patient hypo- and hyperpigmentation of the skin was reported [31]. Loss of *Twist2* function in mice leads to runting, cachexia (adipose and glycogen deficiency), and thin skin with sparse hair [32]. Additional findings were corneal thinning and a reduced population of stromal keratocytes, which are derived from the neural crest [33].

bHLH transcription factors are known to dimerize via their bHLH domain and each monomer binds to one strand of the DNA binding site. MITF is also a bHLH transcription factor absolutely required for the development of melanocytes. TWIST2 has been reported to heterodimerize with other bHLH transcription factors [23]. Thus, we suggest the following mechanism for the expression of the belt phenotype: Ectopically expressed TWIST2 in the developing neural crest of belted cattle might interact in a dominant-negative manner with other bHLH transcription factors (MITF or its partners) and thus lead to either a reduction in the number of cells programmed for the melanoblast lineage or a reduced survival/proliferation of melanoblasts in the developing neural crest. Instead of this protein-protein interaction model with other bHLH transcription factors, ectopically expressed TWIST2 might also directly bind to DNA and repress the transcription of genes essential for melanoblast fate determination, such as e.g. *MITF*.

As a first test of these hypotheses we asked whether overexpressing bovine TWIST2 in zebrafish embryos had an effect on melanocyte development. The TWIST2 protein is highly conserved among vertebrates. Bovine TWIST2 (160 aa) is 100% identical with human TWIST2 and it has 85% identity with zebrafish twist2 (163 aa). We thus hypothesized that overexpression of bovine TWIST2 in the premigratory neural crest of zebrafish might cause a reduction in the number of neural-crest derived melanocytes. Our data indeed indicated that overexpression of TWIST2 led to a decrease in melanocyte numbers in transgenic zebrafish. We were unable to detect EGFP-expressing cells in our TWIST2-overexpressing embryos, suggesting that overexpression was either inconsistent with cell survival or resulted in repression of the *mitfa* promoter. These observations, in conjunction with the pigmentary effects of mutant alleles in humans, are consistent with our proposal that mis-regulated expression of TWIST2 results in the belted pigmentation phenotype. A definitive test will require access to bovine embryos in belted and non-belted animals.

Taken together, the genetic data strongly support a causal role of the CNV in the 5’-flanking region of the *TWIST2* gene for the belted phenotype in cattle. Our functional data further support the hypothesis of a regulatory variant that leads to ectopic expression of TWIST2, which decreases the number of melanocytes in the developing embryo. In common with other mammals, the midsection of the trunk is particularly sensitive to such a decrease, resulting in a characteristic belt phenotype. Belted cattle thus probably represent a spontaneous mutant, in which a structural variant led to subtle changes in gene expression and consequently to an appealing coat color phenotype that was actively selected by cattle breeders. This study highlights the scientific value of spontaneous mutants, which have arisen during animal domestication. Billions of domestic animals have been handled and observed by their human owners providing us with all kinds of fascinating heritable phenotypes. Given the essential role of TWIST2 in development and the dramatic consequences of *TWIST2* coding variants, it is not surprising that the specific mechanism leading to the bovine belt phenotype so far has only been observed once in the entire history of animal domestication.

## Materials and Methods

### Ethics statement

All animal experiments were performed according to the local regulations. The collection of cattle samples was approved by the ‘‘Cantonal Committee For Animal Experiments’’ (Canton of Bern; permit BE75/16). Zebrafish were raised at the Institute of Anatomy, Univesity of Bern (animal house licence number BE413 and animal experimentation number BE15/95. Zebrafish experiments at the University of Bath were reviewed by the Animal Welfare and Ethical Review Body (AWERB), and covered under the Home Office Project License PPL30/2937.

### Cattle samples

We isolated genomic DNA from hair roots or EDTA blood samples from 116 belted animals (6 belted Brown Swiss, 72 Belted Galloway, 38 Lakenvelder) and 264 non-belted controls from different breeds for SNP chip genotyping and the fine-mapping experiment. For the subsequent association study we recruited more than 1,000 additional DNA samples from the routine parentage testing performed at the University of Göttingen. The phenotype of the additional animals was solely based upon owners’ reports. During the investigation we excluded several animals that were reported as belted by the owners, but had only minimal white spots that did not resemble the typical belt phenotype. We also excluded several animals that were reported as non-belted by the owners, but turned out be White Galloways, on which a belt would not have been visible [34]. The final cohort thus consisted of 239 belted animals and 1,303 non-belted cattle. In this cohort, we had 2 supposedly belted and 2 supposedly non-belted animals with discordant genotypes (Table 1). We were not able to verify the phenotypes of these four questionable animals as either the owners could not be reached or the animals had been slaughtered several years before the investigation and no photos or other information was available.

### SNP chip genotyping and haplotype analyses

Genotyping for the fine-mapping experiment was done on Illumina bovine HD beadchips containing 777,962 SNPs by GeneSeek/Neogen. Genotypes were stored in a BC/Gene database version 3.5 (BC/Platforms) and phased with SHAPEIT [35]. The haplotypes for chromosome 3 were exported into an Excel-file and visually inspected for shared segments among the 116 cases.

### Whole genome sequencing

We prepared a fragment library with approximately 300 bp insert size from a homozygous belted Brown Swiss cow (GU21) and a homozygous Belted Galloway bull (BG011) and collected 2 x100 bp paired-end reads on an illumina HiSeq2000 instrument. We obtained 16.6x coverage on GU21 and 8.6x coverage on BG011. We mapped the reads to the UMD 3.1 reference genome with the Burrows-Wheeler Aligner (BWA) version 0.5.9-r16 with default settings [36]. After sorting the mapped reads by the coordinates of the sequence with Picard tools, we labeled the PCR duplicates also with Picard tools (http://sourceforge.net/projects/picard/). We used the Genome Analysis Tool Kit (GATK version 0591) to perform local realignment and to produce a cleaned BAM file [37]. Variant calls were then made with the unified genotyper module of GATK. For variant calling we used only reads with mapping quality of ≥ 30 and bases with quality values ≥ 20. The variant data output file obtained in VCF format 4.0 was filtered for high quality SNPs using the variant filtering module of GATK. The hard filtering of variants was done as explained in the GATK best practice manual. The snpEFF software [38] together with the UMD 3.1 annotation (ENSEMBL release 79) was used to predict the functional effects of detected variants.

### Droplet digital PCR (ddPCR)

To determine the copy number of the CNV on chromosome 3 we designed a ddPCR assay. Primers and probes were designed using the Primer3 program (http://bioinfo.ut.ee/primer3-0.4.0/primer3/). An 86 bp amplicon spanning chr3:118,612,280-118,612,365 was used (fwd GATGAGTGTTCTGGGTGGAAG; rev CTGTGTCTGCCCATCTCTG; probe FAM-AGGTCCCTAGTCTCTGCCTTCCC-BHQ1) to detect the CNV. A 96 bp fragment of the coagulation factor II thrombin (F2) gene (chr15:77,533,575-77,533,670) served as control amplicon. This genomic region showed equal copy-numbers among the sequenced cattle genomes and has no known role in coat color genetics. The sequences of the primers and probe for the *F2* gene were: fwd CCTGTCTGCTGAGACGCCG; rev GTGGTAGAGTTGATTCTGGAATAGAAAGCAT; probe HEX-CCCCGCCACCCGCAGTGTCT-BHQ1. Between 50-150 ng of genomic DNA were digested using 10 units of *Bam*HI restriction enzyme. The CNV has a *Bam*HI restriction site at position chr3:118,612,241, close to the ddPCR amplicon. The restriction digest was prepared in 1x ddPCR Supermix for Probes (Bio-Rad). After incubation at 37°C for 60 min, primers (900 nM each) and probes (250 nM each) were added directly to the mix and droplets were generated using the QX200 Droplet Generator (Bio-Rad). The reactions were amplified with the following temperature profile: initial denaturation for 10 min at 95°C, followed by 40 cycles of 30 s at 95°C and 60 s at 55°C, and a final droplet stabilization at 98°C for 10 min. Droplets were analyzed with the QX200 Droplet Reader and the software provided by the manufacturer (Bio-Rad).

### RNA-seq

We collected 6 mm skin biopsies from 4 adult belted cows (BG0155, GU0448, GU0466, GU0469) and 3 adult non-belted cows (GU0449, GU0467, GU0468) and isolated total RNA. The TruSeq stranded mRNA library prep kit (Illumina) was used to prepare libraries, which were sequenced on the Illumina HiSeq 3000 platform using 2 x 150 bp paired-end sequencing cycles. The reads in FASTQ format were processed for quality checks using the FastQC tool (v0.10.1; http://www.bioinformatics.babraham.ac.uk). Good quality reads were then mapped to the cattle reference genome UMD 3.1 using the STAR Aligner [39]. We allowed for a maximum of four mismatches per read pair and a maximum intron length of 100 kb. We used htseq-count from the HTSeq package to count reads against annotated Ensembl genes models (version 79) Differential expression was carried out on raw read counts for genes using edgeR [40].

### Recombinant constructs for the expression of bovine TWIST2

An 4.1 kb *mitfa* promoter fragment, the coding sequences for EGFP and bovine TWIST2, and the IRES casette were amplified separately and engineered into a vector containing Tol2 sites (pTKXIGΔin; Kawakami Laboratory) using the Gibson Assembly® Cloning kit from NEB. We thus generated the plasmids pmitfa_EGFP (control construct without TWIST2 expression cassette) and pmitfa_btaTWIST2_EGFP. A detailed map of these plasmids is given in Figures S2 and S3. Plasmid DNA was purified using the Qiagen HiSpeed Maxiprep kit and the correct sequence was verified by Sanger sequencing.

### Zebrafish embryo microinjections

Zebrafish were raised according to Swiss regulations at the Institute of Anatomy, University of Bern (animal house licence number BE413 and animal experimentation number BE15/95 from the Ethical Commitee of the Canton Bern). Injections were performed on embryos of wildtype strains AB and Tg(myl7:GFP) [41] at the one cell stage. Approximately 2 to 3 nl of injection mixture was injected in each embryo using a FemtoJet Microinjector (Eppendorf). Injection mixture comprised 1 µl of construct DNA at 100 ng/µl concentration, 1 µl of Tol2 transposase mRNA at 50 ng/µl and 1 µl of phenol red for identification of injected embryos. Six to eight hours after the injections embryos were sorted and unfertilized or damaged embryos were discarded.

### Melanocyte imaging and counting

At approximately 24 hpf, the dishes were cleaned and the medium was changed. The embryos were grown at 28°C and screened at about 35 hpf. Both the controls and TWIST2 expressing embryos were carefully dechorionated, put in tricaine treated fish water (final tricaine concentration 160 mg/l) and used for melanocyte counting. For counting purposes, the embryos were imaged using bright field technique under an SMZ25 fluorescent stereoscope (Nikon). The embryos were imaged lying laterally with their heads facing to the right using methyl cellulose to prevent them from drying. The images were coded and melanocytes were counted by an investigator who was blinded to the information whether embryos were injected with the control construct pmitfa_EGFP or with pmitfa_btaTWIST2_EGFP.

### Antibody labelling and fluorescent microscopy

After counting melanocytes, the embryos were fixed with 2% PFA diluted in PBS, overnight at 4°C and were processed for antibody labelling as described previously [42]. The anti-GFP antibody was obtained from Aves Labs Inc., the anti-Sox10 antibody was obtained from Gene tex. Anti-GFP and anti-Sox10 were used in a dilution of 1:500. Secondary antibodies were anti-chicken coupled to Alexa Fluor® 568 (1:500; ThermoScientific) and anti-rabbit coupled to Alexa Fluor® 647 (1:250; Invitrogen). Nuclei were counterstained with 4c,6-diamidino-2- phenylindole dihydrochlorid (DAPI; Invitrogen). After antibody labelling, the embryos were imaged under an AxioLSM880 confocal microscope (Zeiss). Whole mount embryos were imaged under 10x, the images were processed using ImageJ and Adobe Illustrator.

### Sequence data accession numbers

Whole genome sequencing data and RNA-seq data from this study have been submitted to the European Nucleotide Archive under accessions PRJEB14604 and PRJEB14606.

## Acknowledgements

We thank all owners and breeders who provided samples and information on their animals. The following people are especially acknowledged for contributing to this study: Michèle Ackermann, Nathalie Besuchet-Schmutz, Bernard Conrad, Joëlle Dietrich, Muriel Fragnière, Anne Gesell, Michael Hässig, Javier Langa, Inês Marques, Andrea Patrignani, Marcos Sande, Nathalie Schuster, and Brigite Simoes Rodrigues. We thank the Next Generation Sequencing Platform of the University of Bern and the Functional Genomics Center Zurich for performing experiments. Computationally intensive tasks were partly performed at the Vital-IT high-performance computing centre of the Swiss Institute of Bioinformatics (http://www.vital-it.ch/).

## Supporting Information

**Figure S1.** Historical evidence for the origin of the belted phenotype.

**Figure S2.** Map of expression plasmid pmitfa_EGFP.

**Figure S3.** Map of expression plasmid pmitfa_btaTWIST2_EGFP.

**Table S1.** Fine-mapping of the belt locus.

**Table S2.** RNA-seq experiment.

